# Embryonic signals perpetuate polar-like trophoblast stem cells and pattern the blastocyst axis

**DOI:** 10.1101/510362

**Authors:** Javier Frias-Aldeguer, Maarten Kip, Judith Vivié, Linfeng Li, Anna Alemany, Jeroen Korving, Frank Darmis, Alexander van Oudenaarden, Niels Geijsen, Nicolas C. Rivron

## Abstract

The early mammalian conceptus (blastocyst) comprises an outer trophoblast globe that forms an axis originating from the inner embryonic cells. From the mouse conceptus, Trophoblast stem cells (TSCs) are derived, which are in vitro analogues of early trophoblasts. Here, we show that TSCs contain plastic subpopulations reflecting developmental states ranging from pre- to post-implantation trophoblasts. However, upon exposure to a specific combination of embryonic inductive signals, TSCs globally acquire properties of pre-implantation polar trophoblasts (gene expression, self-renewal) juxtaposing the inner embryonic cells, and an enhanced, homogeneous epithelial phenotype. These lines of polar-like TSCs (pTSCs) represent a transcriptionally earlier state that more efficiently forms blastoids, whose inner embryonic cells then induce the patterning of gene expression along the embryonic-abembryonic axis. Altogether, delineating the requirements and properties of polar trophoblasts and blastocyst axis formation in vitro provides a foundation for the precise description and dissection of early development.

## Introduction

Once the mouse conceptus has completed 4 rounds of cell divisions (day 3 -3,25), both Inner Cell Mass (ICM) and Trophectoderm (TE) tissues arise as the first precursors of the embryo and placenta, respectively (Cross et al., 2003). These two tissues are permissive for the derivation and culture of lines of Embryonic Stem Cells (ESCs) and Trophoblast Stem Cells (TSCs), which are *in vitro* analogues retaining many of the properties of their parental tissues (Roberts et al., 2011; Rossant, 2001).

Both ESCs (Ying et al., 2008) and TSCs (Kuales et al., 2015; Motomura et al., 2016) display a degree of inter-cellular heterogeneity in culture, which reflects the concomitance of different peri-implantation stages. This heterogeneity is facilitated by the use of sub-optimal culture conditions, often not chemically-defined, and that lack the specific combination of signaling molecules able to capture one constrained state. Although inter-cellular heterogeneity represents an obstacle in the establishment of reliable and consistent *in vitro* models, it also allows studying the mechanisms regulating these developmental transitions. Indeed, the heterogeneity of ESCs is reduced by activating Wnt and inhibiting ERK signaling pathways (Ying et al., 2008), thereby constraining the cells in a pre-implantation epiblast-like state. In contrast, despite recent improvement in the culture of TSCs (Erlebacher et al., 2004; Kubaczka et al., 2014; Ohinata and Tsukiyama, 2014), current culture conditions are permissive for a notorious yet ununderstood heterogeneity (Kuales et al., 2015; Motomura et al., 2016).

Intriguingly, lines of TSCs can be derived from both pre-implantation and post-implantation stages (Roberts et al., 2011; Tanaka et al., 1998), which argues for a heterogeneity, plasticity or interconvertibility of trophoblast states. Delineating their developmental equivalence has been challenging but, lately, single cell transcriptomics opened new capabilities. In the conceptus, these states are partly constrained by the local environment, including molecular signals arising from neighbors (e.g. embryonic inductions). Such molecules act in concert and in transient combinations that are difficult to systematically explore *in vitro*.

Here, following an unbiased identification of TSCs subpopulations *via* single cell transcriptomic, we pinpointed multiple convertible developmental states, whose progression is driven by the spatial location within colonies. Then, through systematic combinatorials using factorial design, we exposed a cocktail of signals present in the conceptus that restrain TSCs into a pre-implantation state.

Critically, we show that these inductions are sufficient to spontaneously and spatially pattern gene expression along the embryonic-abembryonic axis (Rossant, 2004; Rossant and Tam, 2009).

## Results

### The transcriptome of polar and mural trophectoderm cells

In the blastocyst, trophoblasts range from a proliferative progenitor state juxtaposing the ICM (polar trophoblasts), to a more differentiated state opposite (mural trophoblasts)(Cross et al., 2003; Gardner, 2000; Rossant and Tam, 2009). We observed that, as blastocysts progress, the expression of the core transcription factor Cdx2 (Niwa et al., 2005; Strumpf et al., 2005) decreases in the mural trophoblasts at both the mRNA (Figure 1A) and protein (Figure 1B, S1A) levels. To determine what other genes are differentially expressed between along the axis, we performed transcriptome analysis on a dataset of single trophoblasts dissected from E4.5 blastocysts (Nakamura et al., 2015). We found 989 differentially expressed genes (FC > 1.5, adj p-value < 0.001, Figure S1B, Table S1). Polar cells were characterized by an increase of regulators of self-renewal (*Cdx2, Esrrb (Gao et al., 2018; Latos et al., 2015a; Luo et al., 1997; Niwa et al., 2005; Rayon et al., 2014; Strumpf et al., 2005)*, proliferation (*Mki67, Cdk1, Ccnd1-3*), and other markers (e.g. *Ddah1* and *Ly6a*). In contrast, mural cells more highly expressed differentiation genes including *Ascl2* (Guillemot et al., 1994), *Ndrg1 (Fu et al., 2017), Gata2 (Ma et al., 1997), Slc2a3 (*a.k.a *Glut3) (Schmidt et al., 2009)*(Figure 1D and Sup table 1, (Gao et al., 2018; Latos et al., 2015a; Luo et al., 1997; Niwa et al., 2005; Rayon et al., 2014; Strumpf et al., 2005)).

**Figure 1.**
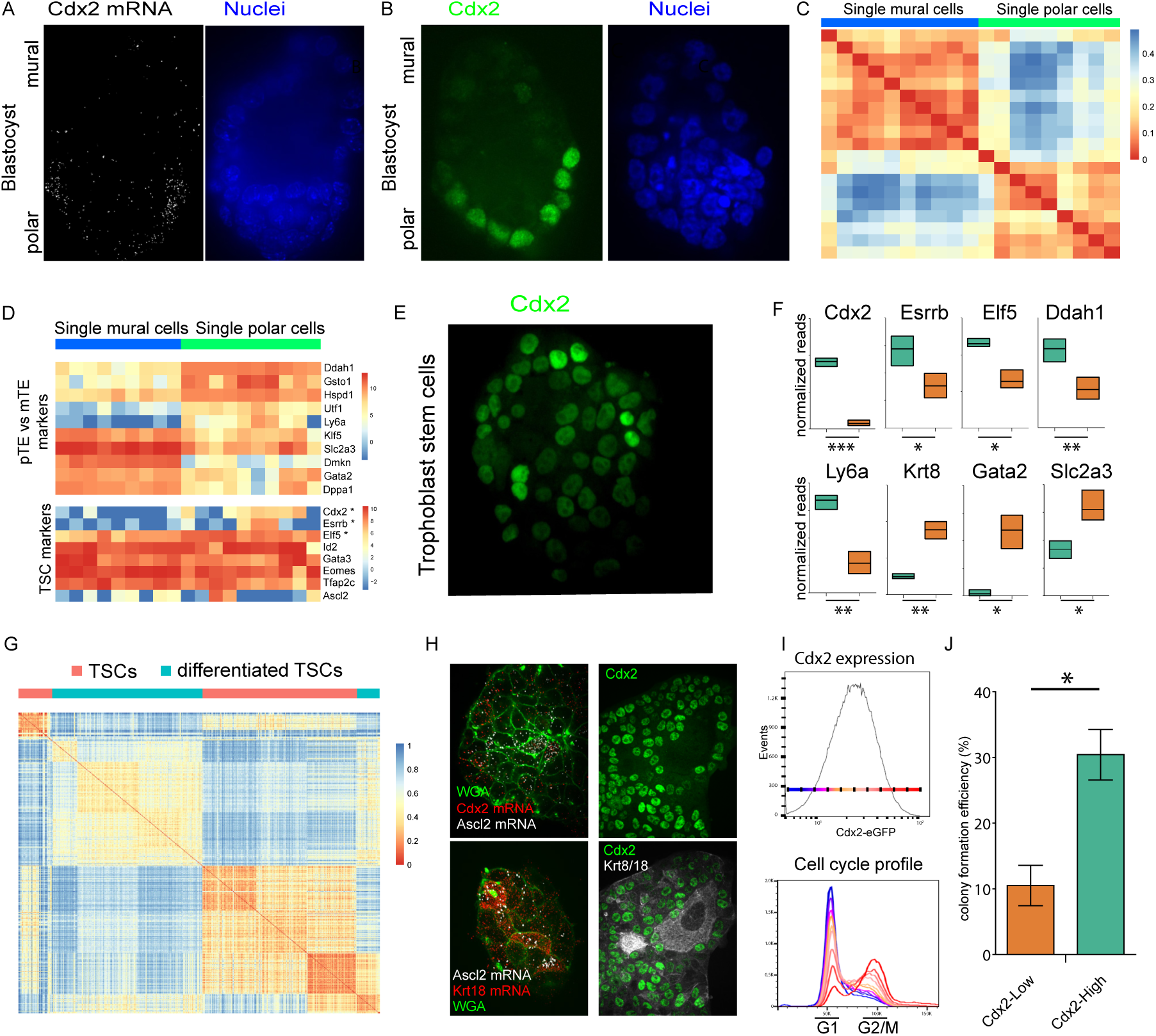
Trophoblast stem cells are heterogeneous and reflect multiple pre- and post-implantation developmental states. A. smFISH for Cdx2 mRNA on E4.5 blastocysts. Nuclei are stained with Hoechst. B. CDX2 protein detection in E3.75 blastocyst after being cultured for 6 hours in plain M2 media. C. Heterogeneous expression of CDX2 in cultured TSCs (also see I). D. Unsupervised gene clustering analysis of E4.5 trophectoderm single cells (Nakamura et al.). E. Gene expression heatmap for single TE cells. F. Differential gene expression of differentiation markers in CDX2-High (green) and CDX2-Low (orange) populations. * for p value<0.05, ** for p value< 0.01 and *** for p value<0.001. G. Unsupervised cell clustering analysis of TSCs and 6-days differentiated TSCs. H. smFISH for Cdx2 and Ascl2 (post-implantation ExE marker (Guillemot et al., 1994)) transcripts in TSCs (top left) or for Ascl2 and Krt18 (bottom left). Anticorrelation for CDX2 and Krt8/18 expression in TSCs as assessed by immunofluorescence (right). I. Cell cycle analysis of TSCs cultures shows a correlation between CDX2 expression (higher CDX2 content in red, lower in blue) and the cell cycle state. J. Colony formation potential of single CDX2-High and CDX2-Low cells. fc=2.89, pvalue=0.015.

### TSCs are heterogeneous and mirror multiple developmental states

We wished to determine whether TSCs, similar to TE cells, showed heterogeneity in the expression of Cdx2 and other genes. Indeed, TSCs cultured *in vitro* displayed intercellular heterogeneity for Cdx2 (Figure 1E, Cdx2-eGFP TSCs (McDole and Zheng, 2012)) as also noted previously (Kuales et al., 2015; Motomura et al., 2016). We thus analysed the transcriptome of Cdx2-High and Cdx2-Low TSCs (1000 cells sorted from top and bottom 5%, respectively, of Cdx2-eGFP expressing cells). By this comparison, 1941 genes were differentially expressed between the two populations (FC > 1.5, p-value < 0.01, Table S2). Cdx2-High TSCs were enriched for transcripts regulating trophoblasts self-renewal (*Cdx2, Esrrb* (Gao et al., 2018; Latos et al., 2015a; Strumpf et al., 2005)), the cell cycle, which is consistent with polar trophoblasts (Gardner, 2000), the Hippo (Menchero et al., 2017; Nishioka et al., 2008, 2009; Posfai et al., 2017; Rayon et al., 2014; Yagi et al., 2007) and Tgfb (Erlebacher et al., 2004; Kubaczka et al., 2014) pathways regulating trophectoderm development (Figure S1C, Table S3), and polar transcripts such as *Ddah1* and *Ly6a*. In contrast, the Cdx2-Low population showed higher expression of differentiation markers including *Ndrg1, Gata2, Slc2a3* (a.k.a. *Glut3*), a nd *Krt8* (Figure 1F)(Nakamura et al., 2015; Nishioka et al., 2008; Ralston et al., 2010; Rivron et al., 2018b). Together, these data indicate that the transcriptome of Cdx2-High TSCs resembles the one of polar trophoblasts, while Cdx2-Low cells show signs of developmental progression.

In ESCs, the subpopulations reflect closely related developmental states that are convertible (Godwin et al., 2017; Hastreiter and Schroeder, 2016; Luo et al., 2013; Ochiai et al., 2014; Ying et al., 2008). To determine whether the subpopulations of TSCs are convertible in culture, we sorted Cdx2-High and Cdx2-Low TSCs and cultured them separately. Within 5 days, the heterogeneity was re-established in all conditions (Figure S1D). We concluded that at least part of the Cdx2-high and Cdx2-low subpopulations are interconvertible, and that intercellular heterogeneity is an intrinsic property maintaining an equilibrium of subpopulations under these culture conditions.

### Subpopulations of TSCs reflect both blastocyst and post-implantation stages

To unbiasedly delineate the subpopulations of TSCs, we performed single cell transcriptome analysis on 330 TSCs and 332 differentiated TSCs (cultured for 6 days without growth factors). Unsupervised cell clustering analysis separated both cultures and their subpopulations (Figure 1G). Interestingly, one subpopulation of TSCs clustered with the differentiated TSCs, suggesting that a fraction of the cells present in the TSC culture is differentiated. Monocle analysis (Trapnell et al., 2014) arranged all single trophoblasts in a pseudotime trajectory containing 6 clusters (4 clusters for TSCs and 2 clusters for differentiated TSCs, Figure S1E–F). Low pseudotime values corresponded to a subpopulation of TSCs with higher expression levels of *Cdx2, Esrrb, Eomes* (Figure S1G). Cells with intermediate pseudotime values showed higher expression of the extraembryonic ectoderm (ExE) markers *Elf5* (Donnison et al., 2005; Latos et al., 2015b; Ng et al., 2008) or *Ascl2 (Guillemot et al., 1994)*. High pseudotime values corresponded to cells expressing terminal differentiation markers such as *Flt1* or *Gcm1* (Figure S1G)(Baczyk et al., 2004; Hughes et al., 2004; Simmons and Cross, 2005). Again, a subpopulation of TSCs (9,2%) was assigned the high-pseudotime values of differentiated trophoblasts. We extracted the main genes defining such trajectory, thus unbiasedly revealing markers of the multiple states (Figure S1H). These results show that TSCs include multiple subpopulations reflecting a developmental progression and including differentiated trophoblasts (∼10%).

Single molecule FISH against *Cdx2* and *Ascl2* mRNAs in TSCs colonies confirmed that cells expressing high levels of *Cdx2* transcripts simultaneously expressed low levels of *Ascl2* mRNA (Figure 1H, top left) and low levels of *Krt18* (Figure 1H, bottom left). Accordingly, high protein levels of Cdx2 correlated with low protein levels of Krt8 and Krt18 (Figure 1H, right and bottom right). The differentiated *Ascl2+* and *Krt18+* cells were mainly located in the central part of large colonies, while cells on the edges were more abundant in *Cdx2* transcripts.

Altogether, we concluded that the culture of TSCs is permissive for the concomitance of convertible subpopulations reflecting the polar TE (*Cdx2*^*high*^, *Esrrb*^*high*^, *Ddah1+, Ly6a+*) and mural and post-implantation trophoblasts (*Cdx2*^*low*^, *Esrrb*^*low*^, *Gata2+, Ascl2+*). The spatial location of cells within colonies contributes to the developmental progression, with outer cells more prominently expressing *Cdx2* and the inner cells more prominently expressing *Ascl2* (Figure S1E and Fig 1H).

### The Cdx2-High cells have proliferative and self-renewing features of polar trophoblasts

We next examined the functional differences between Cdx2-High and Cdx2-Low populations of TSCs. The Cdx2-High cells were present in all phases of the cell cycle, while the Cdx2-Low cells were predominantly in the G0/G1 phases (Figure 1I). We also tested the potential for self-renewal by examining the clonogenicity of isolated TSCs. Single Cdx2-High cells formed 3 times more colonies (30% of single cells) as compared to Cdx2-Low cells (10% of single cells, Figure 1J). These observations show that Cdx2-High TSCs have higher proliferative and self-renewing capabilities. The higher clonogenic capability of Cdx2-high cells is consistent with the more proliferative and self-renewing state of polar trophoblasts sustaining the development of the placenta (Gardner, 2000). Although tempered, the capacity of Cdx2-low cells to form colonies is consistent with our observation of convertible states (Figure S1D). Altogether, we concluded that TSCs are heterogeneous and include convertible, closely related developmental states of both pre-implantation and post-implantation trophoblasts.

### Combined embryonic signals generate Cdx2-High trophoblast stem cells

Previous studies demonstrated that inductive signals originating from the inner embryonic cells are necessary to sustain trophoblast proliferation and self-renewal (Gardner, 2000; Guzman-Ayala et al., 2004; Rivron et al., 2018b; Tanaka et al., 1998). We thus reasoned that a combination of these signals could reduce the heterogeneity of TSCs and coerce a phenotype comparable to polar trophoblasts. We formed a library of compounds (Table S4) based on the pathways active in the trophectoderm (Figure S2A) and on ligands present in the blastocyst (Figure S2B). We initially tested these compounds individually for their capacity to regulate Cdx2-eGFP expression in chemically defined medium and on Matrigel-coated plates (Figure 2A and Figure S2C)(Kubaczka et al., 2014)). We found 9 compounds (Il11, 8Br-cAMP, activin, XAV939, Bmp4, Bmp7, LPA, retinoic acid and rosiglitazone) positively regulating Cdx2 expression in a dose-dependent fashion (Figure S2D, Cdx2-High/Cdx2-LOW ratio > 1.25).

**Figure 2.**
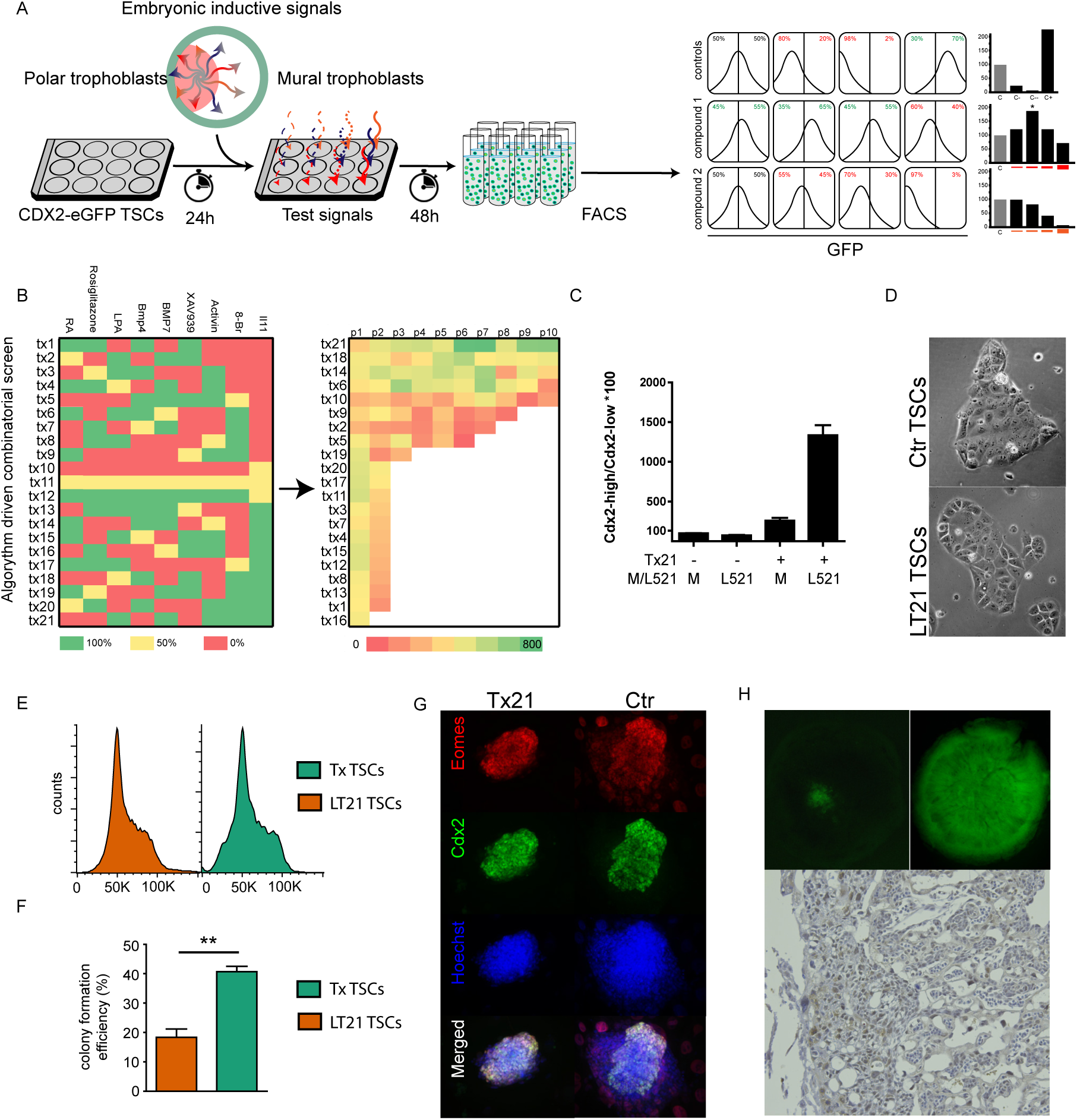
Combined embryonic inductive signals coerce a potent CDX2-high trophoblast state. A. Strategy followed in order to identify CDX2 expression regulators in CDX2-eGFP TSCs. B. List of 21 different cocktails combining the 9 different CDX2 regulators (left). The 21 cocktails are ranked based on the average CDX2-High/ CDX2-Low ratio (right). C. Comparative CDX2-High/CDX2-Low ratio of TSCs upon combination of the top cocktail (Tx21) with Laminin 521. D. Bright field microscopy pictures of TSCs colonies cultured in Tx or LT21 culture conditions. E. Cell cycle profile of TSCs (orange) and LT21 TSCs (green). F. Colony formation efficiency of single sorted control (orange) or LT21 TSCs (green). G. CDX2 and EOMES staining of blastocyst outgrowths 4 days after plating. H. Detection of GFP signals in E10.5 placentas obtained upon the injection of blastocysts with LT21-converted TSCs (top left) and LT21-derived TSCs (top right). Immunostaining against GFP on tissue sections obtained from chimeric placentas (bottom).

Because these signals are likely to act synergistically in the blastocyst, we used factorial design to systematically explore the combinatorial landscape without performing all combinatorials empirically (Hutchens et al., 2007). We cultured Cdx2-eGFP TSCs for 10 passages in 21 different cocktails selected by factorial design (Figure 2B, left) while monitoring Cdx2 expression at each passage (Figure 2B, right). All cocktails resulted in an initial upregulation of Cdx2 expression; however, most cultures (17/21) collapsed after two passages due to a lack of proliferation or attachment. The most likely explanation is that these combinations lead to trophoblast differentiation. The remaining four conditions led to a sustainable, long-term increase in Cdx2 expression. Principal component analysis identified Il11, 8-Br cAMP, BMP7 and LPA as beneficial for the long-term expression of Cdx2, while XAV939, rosiglitazone and retinoic acid, despite their short-term effect, were detrimental (Figure S2E and S2F).

To further investigate the phenotype exhibited in these new conditions, we analysed the transcriptome of TSCs cultured in the 4 effective cocktails. All conditions retained an undifferentiated TSC identity as denoted by the higher expression of self-renewal markers (*Cdx2, Eomes, Esrrb*) and the lack of upregulation of differentiation markers (Figure S2G). We selected one condition (Tx21) based on the higher expression of *Cdx2* and *Eomes*, and the downregulation of the ExE marker *Ascl2* (Figure S2H). Although not necessarily exhaustive, this exploration of the combinatorial landscape shows that TSCs respond to specific combinations of signals present in the embryo.

In order to obtain fully chemically-defined culture conditions, we then tested 8 different laminin proteins to replace Matrigel coatings (Figure S2I). Three laminins (L221, L511 and L521) resulted in successful TSC cultures, without decreasing proliferation or Cdx2 expression. We selected L521 based on the expression levels in the blastocyst (Figure S2J), and cultured TSCs on L521-coated plates and in Tx21 medium. These culture conditions robustly increased Cdx2 expression and depleted the Cdx2-Low subpopulation as compared to both Matrigel/Tx21 or L521 alone (Figure 2C and Figure S2K). After culture optimization, (data not shown) we found the optimal culture conditions to contain Fgf4 (25 ng/ml), Tgfb1 (2 ng/ml), Activin (50 ng/ml), Il11 (50 ng/ml), BMP7 (25 ng/ml), cAMP (200 nM) and LPA (5 nM) and L521 coating. We termed this new culture condition LT21.

### LT21 TSCs have enhanced self-renewal and contribute to placenta formation

TSCs grown in LT21 culture conditions showed consistent high levels of Cdx2 expression. They also displayed a spatially homogeneous expression of *Cdx2* transcripts across colonies, contrasting with the radial patterns observed in colonies grown in previous culture conditions (Figure S2L). The cell cycle profiles of LT21 and Tx cultures were similar (Figure 2E) but the clonogenic potential of LT21 TSCs was largely enhanced (40% of single cells formed a colony, 2-fold, p-value <0.0028; Figure 2F). This clonogenic potential is now akin to the one of ground-state ESCs (Ying et al., 2008). LT21 TSCs also displayed a cobblestone-like phenotype often observed in epithelial cell types and consistent with the epithelial nature of the trophectoderm (Figure 2D).

LT21 TSCs, conserve their differentiation potential as demonstrated by the rapid decrease in Cdx2 expression (day 1) followed by the upregulation of the ExE marker Ascl2 (peak after three days, Figure S2M) upon growth factor removal (Latos et al., 2015a). LT21 conditions were permissive for the derivation of new TSC lines. Due to a lack of initial attachment and proliferation of blastocysts on L521, the first two passages made use of a layer of mouse embryonic fibroblasts (Figure S2N). Four days after blastocyst plating, the outgrowths in Tx21 medium showed a decreased degree of differentiation (reduction of Cdx2-negative and EOMES-negative cells, Figure 2G) when compared to Tx conditions. Upon further culture, the new established cell lines were tested for *in vivo* potential. Both LT21-converted (originally derived in serum conditions) and LT21-derived TSC lines led to the formation of placenta chimaeras upon injection into foster blastocysts (Figure 2H). Importantly, in previous chemically-defined medium, a transition to serum-containing medium was necessary to generate chimeras (Kubaczka et al., 2014). This was not necessary anymore in LT21 conditions. We thus propose that the LT21 conditions lead to potent TSCs able to rapidly transitions through a post-implantation stage both *in vitro* and *in vivo*.

### The transcriptome of LT21 TSCs reflects the one of polar trophoblasts

We then investigated the balance between LT21 TSC subpopulations *via* single cell transcriptome analysis (LT21 TSCs, TSCs, and differentiated TSCs). LT21 TSCs showed enhanced transcriptional homogeneity as compared to Tx TSCs (unsupervised cell clustering, Figure 3A). Monocle analysis assigned lower pseudotime values to LT21 TSCs as compared to Tx TSCs and to differentiated TSCs (Figure 3B) indicating that LT21 TSCs reflect an earlier developmental state. Low pseudotime and LT21 TSCs were characterized by the expression of the early TE genes *Tead4* (Nishioka et al., 2009; Ralston et al., 2010) and *Klf5 (Lin et al., 2010; Ralston et al., 2010)*, of the TE transcription factors (*Cdx2, Eomes, Esrrb*) as well as polar markers (*Ddah1, Hspd1, Gsto1, Utf1*). This confirmed the directionality of the developmental progression and the earlier identity of LT21 TSCs (Figure 3C). Developmental progression unrolled along the pseudotime trajectory, with the subsequent upregulation of *Elf5 (Hemberger et al., 2010; Ng et al., 2008; Ralston et al., 2010), Tfap2c (Cao et al., 2015; Kuckenberg et al., 2010) Foxd3* (Hanna et al., 2002; Tompers et al., 2005) and *Gata3* (Paul and Home, 2015; Ralston et al., 2010; Ray et al., 2008), which are expressed both at blastocyst and at postimplantation stages and drive self-renewal and differentiation in a context dependent and stoichiometric fashion (Auman et al., 2002; Kidder and Palmer, 2010; Kuckenberg et al., 2010; Ralston et al., 2010). Trophoblasts with higher pseudotime values expressed *Ascl2 (Guillemot et al., 1994), Nodal* (Natale et al., 2009) and *Ets2* (Wen et al., 2007), which marks postimplantation cells in the ectoplacental cone and chorion. These genes are expressed in a subpopulation of TSCs but largely depleted in LT21 TSCs (Figure 3c). Finally, high-pseudotime values were attributed to differentiated trophoblasts expressing *Flt1, Gcm1, Krt8/18, Cited2, Gjb2* and *Nrk (Imakawa et al., 2016; Morioka et al., 2017; Sood et al., 2006a)*(Figure S3D).

**Figure 3.**
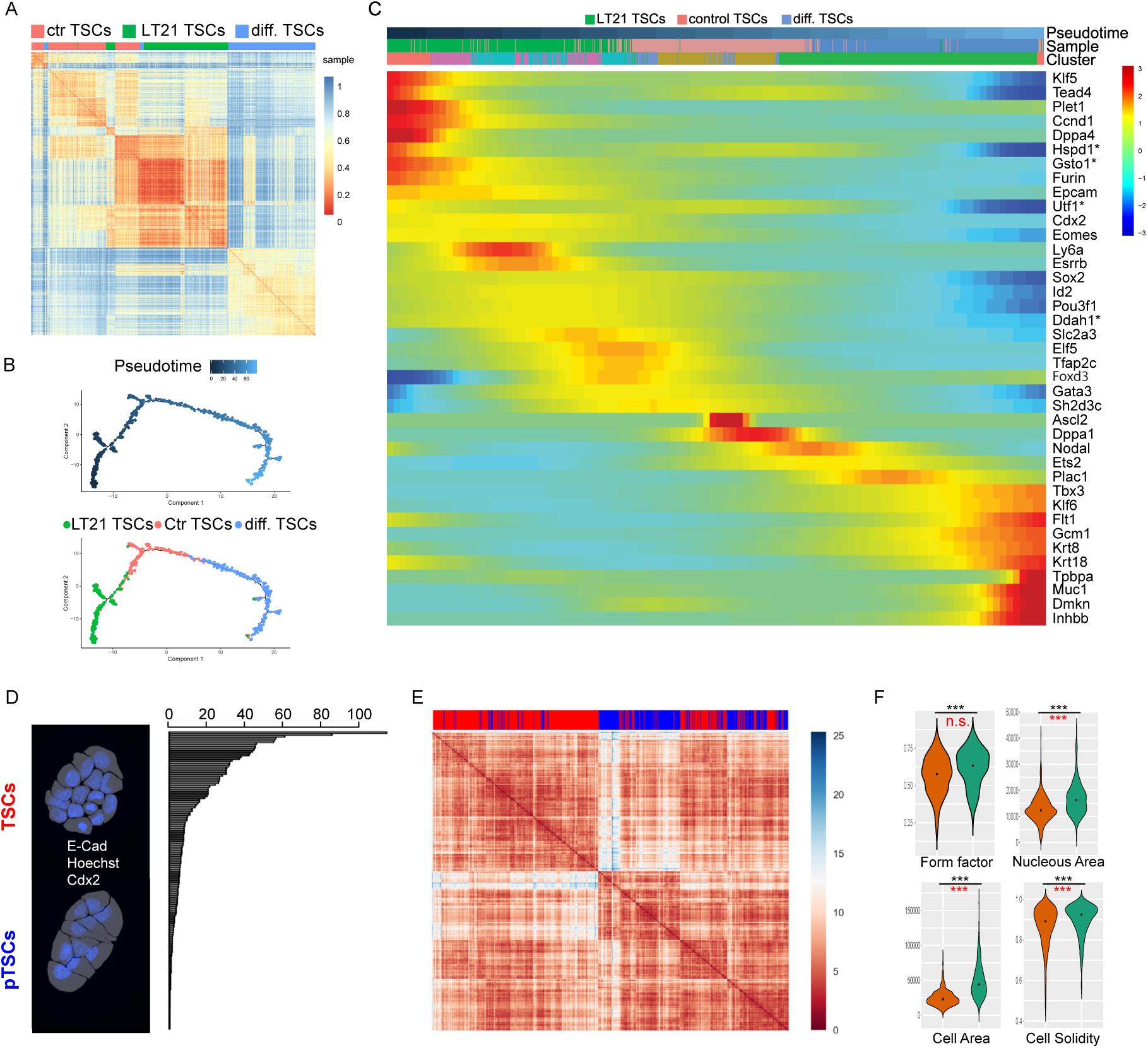
The CDX2-high TSCs reflect an earlier, polar-like state with an enhanced epithelial phenotype. A. Unsupervised gene clustering analyisis of single cells from LT21, control and differentiated TSCs. B. Single cell pseudotime trajectory obtained from LT21, control and differentiated TSCs. C. Pseudotime heatmap for visualization of expression patterns along the pseudotime trajectory along with polar, mural and classical TSC markers. D. High-content imaging and extraction of morphological feature of TSCs and polar-like TSCs (pTSCs) segmented based on E-cadherin, Hoechst, and CDX2 stainings (left). Ranking of differential features based on the p-value after Mann-Whitney statistical analysis (right). E. Heatmap unsupervised clustering based on the top 20% differential features of pTSCs (blue) and TSCs (red). F. Violin plots of the value distribution for some top-ranked differential features. Unpaired two-tailed t-test (black) and F test for variance (red). ***=p-value<0.001.

The unbiased extraction of the 50 genes that better define the pseudotime trajectory (Figure S3D) revealed the expression of genes *Duox2* and *Gsto1* in lower pseudotime, which were previously described as markers for *bona fide* TSCs (Kuales et al., 2015) and polar trophectoderm cells (Nakamura et al., 2015), respectively. We also found that transcripts for epithelial (Epcam, Ecad) and tight junction (*Tjp2, Cldn4, Cldn6*) were more highly expressed in cells with low pseudotime values (Figure S3B). We confirmed *via* qPCR that LT21 TSCs were depleted from differentiated markers (*Ascl2, Gcm1*), while showing higher expression of *Cdx2, Esrrb* and *Ly6a* (Figure S3C). Of note *Ly6a* is a polar marker that was lately proposed to also mark stem cells during placentation (Natale et al., 2017).

Altogether, these results show that LT21 TSCs increase (i) the expression of transcription factors regulating self-renewal (e.g. *Cdx2, Esrrb*), (ii) their functional clonogenicity, (iii) the expression of polar genes (e.g. *Ly6a, Ddah1, Hspd1, Gsto1, Utf1*), while (iv) reducing the intercellular transcriptional variance, (v) depleting the differentiated subpopulations (e.g. expressing *Gmc1, Ascl2, Nodal, Ets2*), and (vi) maintaining the potential to rapidly differentiate and chimerize the placenta. Therefore, we propose to call these cells polar-like TSCs (pTSCs).

### Polar-like TSCs display enhanced epithelial features

The transcriptome of pTSCs showed enrichment in transcripts related to the epithelial phenotype including extracellular matrix organization, cell adhesion, pathways related to ECM-receptor interaction, focal adhesion, cytoskeleton and tight junctions. Especially, genes encoding for the tight junction molecules *Cldn4, Cldn6, Tjp2*, and *Jam2* were upregulated (Figure S3B). Of note, *Cldn4* and *Cldn6* have been shown to be essential for blastocyst formation (Moriwaki et al., 2007). We further investigated these changes by measuring the morphometric characteristics of single cells *via* high-content imaging. E-CADHERIN and Hoechst stainings segmented the plasma membranes and nuclei respectively, which, along with Cdx2 immunofluorescence (Figure 3D), permitted to automatically extract 161 morphometric features from both control TSCs (502 cells) and pTSCs (297 cells). We ranked these features based on the p-value scores (Mann-Whitney, Table S5). The top 20% morphometric features clearly separated the two cell populations (Figure 3E). We concluded that pTSCs are morphologically different from TSCs. Specifically, pTSCs had significantly larger cell and nuclei areas, and were more circular and less lobulated compared to TSCs (Figure 3F). Because TSCs had a similar overall sizes in suspension (Figure S2K), we concluded that pTSCs were more spread than TSCs. Thus, consistent with the higher expression of epithelial genes (*Epcam, Ecad, Cldn4, Cldn6*, Figure S3B), pTSCs have enhanced epithelial morphological features. Finally, we measured the average distance of each cell to the population centroid (100 different tSNE maps), which was smaller for pTSCs compared to TSCs. We concluded that pTSCs were morphometrically more homogeneous compared to TSCs (Figure S3F). Altogether, these data indicate that pTSCs are more morphologically more homogeneous, with an enhanced epithelial-like phenotype (e.g. more spread and less lobulated) and gene expression profile (e.g. *Epcam, Cldn4/6, Jam2, Tjp2*), consistent with the phenotype of TE cells.

### Polar-like TSCs yield higher number, and more circular blastoids

Next, we reasoned that cultures reflecting an earlier developmental state and depleted from post-implantation-like trophoblasts should more efficiently recapitulate features of TE development *in vitro*. We thus tested the potential of pTSCs to form blastoids, a stem cell–based models of the blastocyst (Rivron, 2018; Rivron et al., 2018a, 2018b). pTSCs lines formed blastoids more efficiently than TSCs (Figure 4A). Comparing the morphological features (Figure S4A) of blastoids, we found that pTSCs formed larger, more circular blastoids, and more efficiently formed a cavity compared to TSCs (Figure 4B–C and Figure S4B–D). Next, we tested the intrinsic potential of pTSCs to form trophospheres (formed only with TSCs) and found an improved circularity and cavitation efficiency compared to TSCs-derived trophospheres (Figure S4C, E). We concluded that pTSCs have an enhanced intrinsic potential to recapitulate features of TE development. Because blastocysts and blastoids form through inductive signals originating from the embryonic cells (Gardner, 2000; Rivron et al., 2018b), we looked whether pTSCs better responded to ESCs. pTSCs within blastoids enhanced the diameter of the cyst, a feature that did not occur in trophospheres (Figure S4B). Because pTSCs did not proliferate more than control TSCs (Figure S4F), we concluded that pTSCs better responded to signals responsible for the swelling of the cyst. Blastoids generated from pTSCs correctly localized the basal adherens junctions (E-cadherin)(Larue et al., 1994) and apical cytoskeletal protein (Krt8/18) typical of an epithelium (Figure 4C). Altogether, we concluded that pTSCs have an enhanced potential to model TE features.

**Figure 4.**
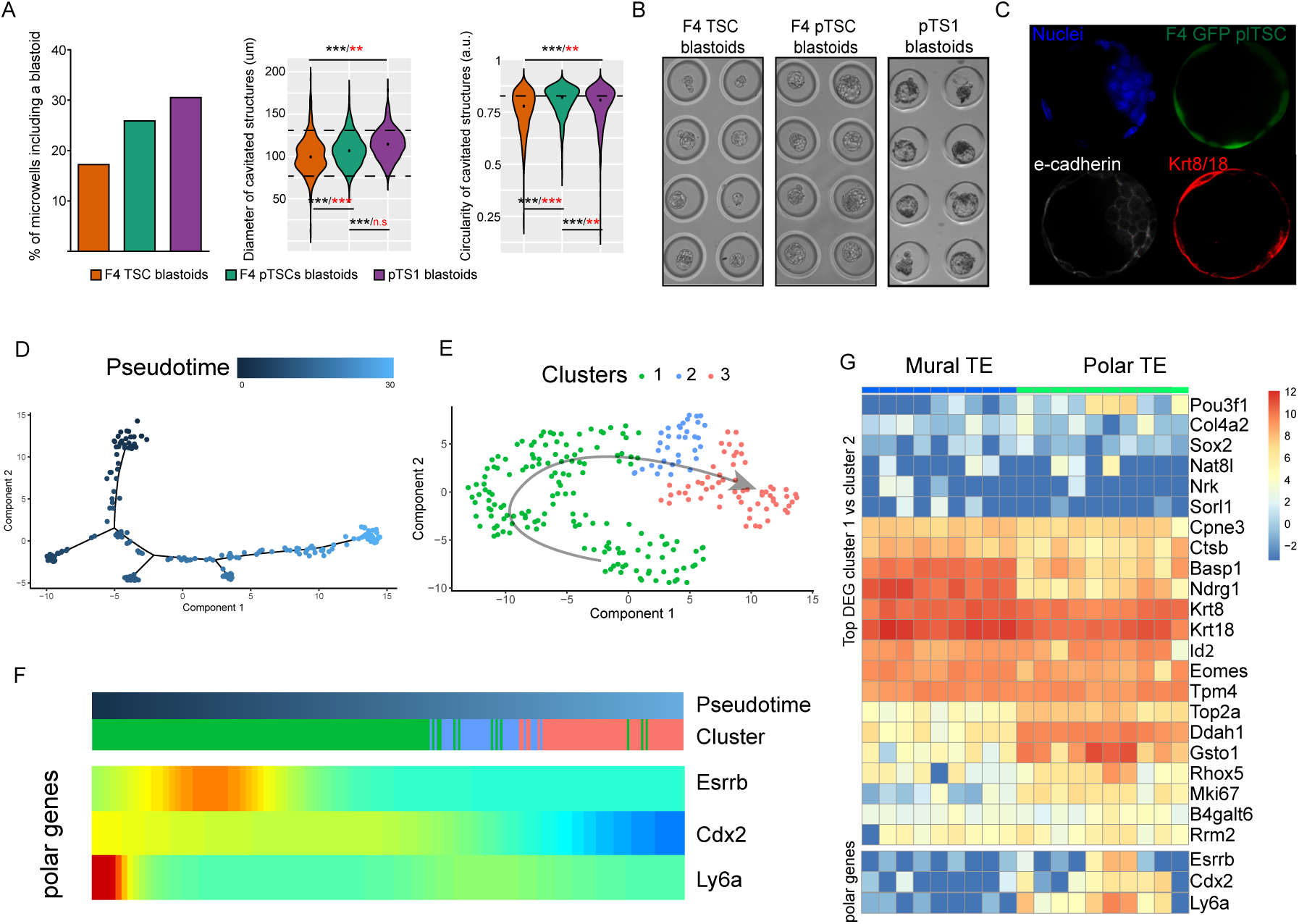
pTSCs efficiently form blastoids and spontaneously pattern gene expression along the embryonic-abembryonic axis. A. Percentage of microwells including a blastoid (based on circularity, diameter and presence of one single cavity) for control TSCs (F4, in orange), pTSCs (F4 converted to LT21, in green), and pTS1 pTSCs (directly derived in LT21 conditions, in purple) (left). Diameter of all structures included in all microwells (center). Circularity of all structures included in all microwells (right). Two-tailed unpaired t-test. ** for p-value<0.01, *** for p-value<0.001. Black represent the analysis of the means. Red represents the analysis for F-variance. B. Representative pictures of microwells including control F4, F4 pTSCs, and pTS1 blastoids. C. Blastoid formed from F4 pTSCs and stained for E-cadherin and KRT8/18. D. Pseudotime trajectory of single trophoblasts isolated from blastoids. E. Clustering analysis from monocle groups the cells in three clusters. F. Pseudotime heatmap for the polar genes Esrrb and Cdx2, and Ly6a. G. The top 25 differentially expressed genes between the polar-like and mural-like clusters plotted for the polar and mural TE cells of E4.5 blastocysts.

### Embryonic inductions contribute to forming the embryonic-abembryonic axis

Finally, we wondered whether the embryonic inductions were sufficient to spatially pattern gene expression along the TE-like cyst of blastoids. We thus isolated single trophoblasts from blastoids and performed single cells sequencing (Figure S4G). As revealed by Monocle, the trophoblasts formed a pseudotime trajectory (Figure 4F) including three transitioning subpopulations (Figure 4G). The cluster with low pseudotime values expressed higher *Cdx2, Eomes, Esrrb* (Table S6 and Figure 4F-G)(Festuccia et al., 2018; Gao et al., 2018; Latos et al., 2015a; Niwa et al., 2005; Russ et al., 2000; Strumpf et al., 2005) along with numerous polar genes (e.g. *Mki67, Ly6a, Gsto1, Ddah1, Utf1, B4galt6*, *Cpne3, Duox2, Nat8l, Pou3f1, Ppp2r2c, Rhox5, Rrm2, Sorl1* and *Top2a*). In contrast, the cluster with high pseudotime values showed higher expression levels of the mural markers *Ndrg1, Basp1, Flt1, Krt8*/*18*,, *Ctsb* and *Slc5a5 (Shi et al., 2013; Sood et al., 2006b)*, similar to the blastocysts (Figure 4G). Using smFISH, we confirmed that *Ly6a* was indeed more prominently expressed in the polar cells in both blastocysts (3/3, Figure S4H) and blastoids (7/10, Figure S4I). We concluded that blastoids recapitulate aspects of the embryonic-abembryonic axis, and that embryonic inductions are sufficient to spatially order gene expression along the trophectoderm (Figure 5).

**Figure 5.**
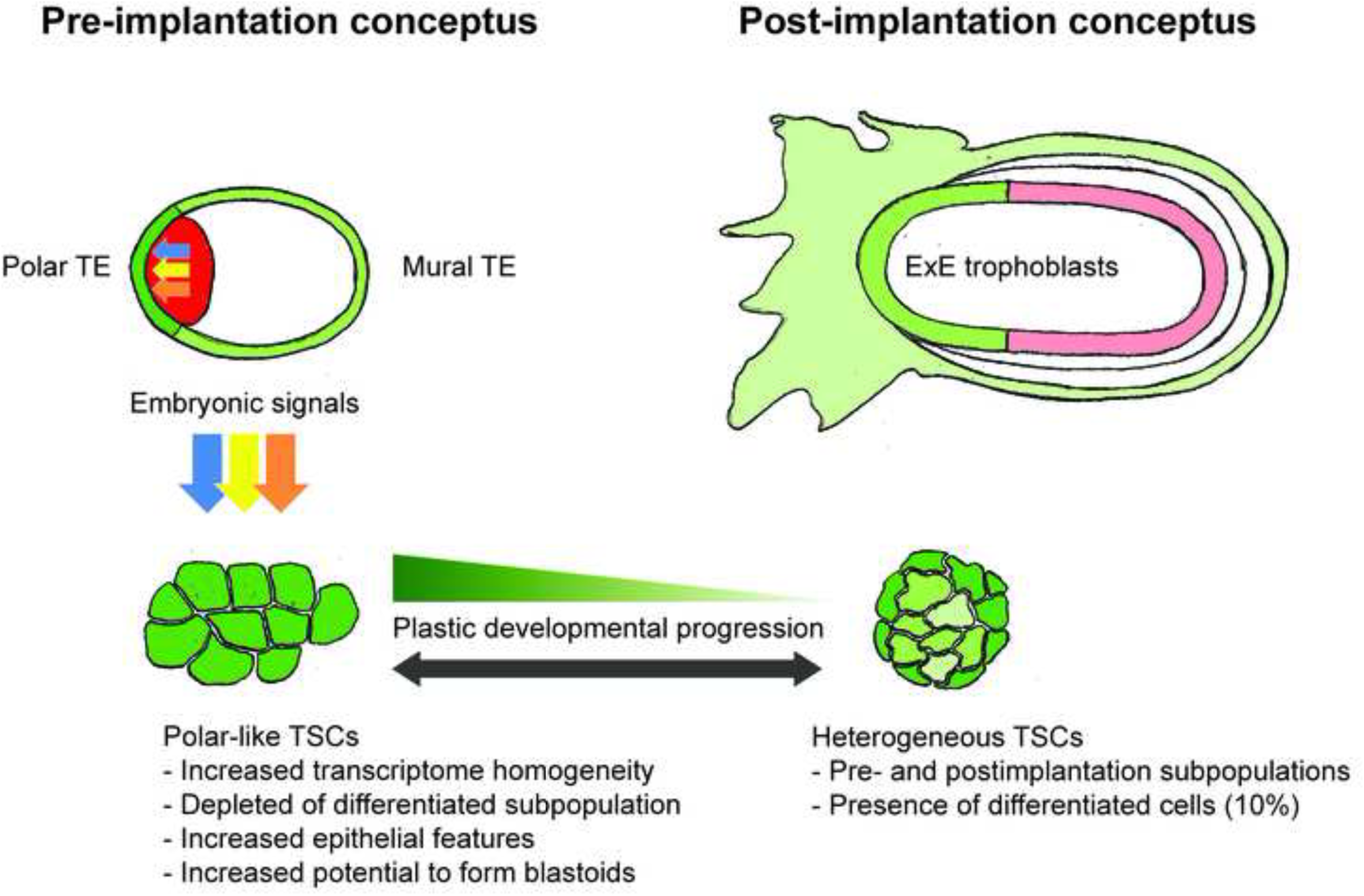

## Discussion

Lines of stem cells are powerful *in vitro* tools to study the nature and functions of the embryonic and extraembryonic lineages. However, the use of sub-optimal culture conditions can prevent capturing constrained cell states, which hampers our understanding of developmental transitions. For example, ESCs cultured in serum-containing medium reflect a range of concomitant peri-implantation states, and the use of chemically defined culture conditions including specific soluble compounds (known as 2i/3i conditions) reduces the degree of intercellular heterogeneity in part by preventing differentiation (Ying et al., 2008). Such culture conditions proved valuable to precisely delineate the mechanisms of pluripotency. It also implied a more efficient derivation of cell lines, and the development of more efficient differentiation protocols (Vrij et al., 2019).

The approach we presented relies on the identification of subpopulations *via* single cell RNA sequencing and on a systematic exploration of combinations of factors present in the blastocyst using factorial design. This approach can potentially be applied to other cell lines (e.g. XEN cells (Niakan et al., 2013)). We defined subpopulations of TSCs reflecting pre-implantation to post-implantation states and including largely differentiated cells (∼10%). The convertibility of some of these subpolations as shown by the recovery of heterogeneity upon FACS sorting and by the clonogenic potential of Cdx2-High and Cdx2-Low cells, is compatible with previous observations that TSCs can be isolated both from the blastocyst and the post-implantation conceptus (Tanaka et al., 1998), and suggests a plasticity of the stem cells present in the trophoblast compartment of the peri-implantation conceptus. Recently, we and colleagues developed *in vitro* models of the early conceptus that recapitulate features of development (Reviewed in (Shahbazi and Zernicka-Goetz, 2018)). These models are useful to dissect biological principles (Rivron et al., 2018a). Building on previous observations (Gardner, 2000), the blastoid, a model of the blastocyst, systematically inventoried inductive signals originating from the embryonic cells, and pinpointed at some of their functions on trophoblast proliferation, self-renewal, and epithelial maturation (Rivron et al., 2018b). Here, we used this inventory to create a library and delineate a combination of signals that constrain TSCs into a state with enhanced self-renewal, homogeneous epithelial-like phenotype, and restrained gene expression, all features consistent with the polar trophoblasts.

These pTSCs efficiently form blastoids that spontaneously generate the gene expression patterns observed along the embryonic-abembryonic axis. We concluded that, following subtle early patterning events (Graham and Zernicka-Goetz, 2016; Zhang and Hiiragi, 2018), the inductive signals originating from the embryonic cells significantly contribute to spatially patterning gene expression along the first axis of the conceptus.

## Experimental procedures

### Cell culture

TSCs were cultured under Tx conditions followed a previously published protocol (Kubaczka et al., 2014). After coating with Matrigel, cells were cultured in Tx media, which consists of phenol red–free DMEM/F12 supplemented (phenol red-free, with l-glutamine), insulin (19.4 μg/ml), l-ascorbic-acid-2-phosphate (64 μg/ml), sodium selenite (14 ng/ ml), insulin (19.4 μg/ml), sodium bicarbonate (543 μg/ml), holo-transferrin (10.7 μg/ml), penicillin streptomycin, FGF4 (25 ng/ml), TGFβ1 (2 ng/ml) and heparin (1 μg/ml). LT21 cultured cells were plated in laminin 521–coated plates (10 ug/ml diluted in PBS with Mg2+ and Ca2+) using Tx media supplemented with Il11 (50 ng/ml), Activin (50 ng/ml), Bmp7 (25 ng/ml), LPA (5 nM) and 8Br-cAMP (200 nM). TSCs were differentiated by refreshing with plain TX media without Fgf4, Tgfb1 nor any other Cdx2 regulator. This media was maintained for 6 days. ESC were cultured under 2i conditions (Ying et al., 2008) in gelatin-coated plates. Cells were routinely passaged using trypsin and quenched with trypsin soybean inhibitor.

### TSC line derivation

E3.5 blastocysts were isolated from pregnant females. Zona pellucida was removed using Tyrode acid solution and then they were placed in MEF-coated plates in the presence of Tx or Tx21 media. Media was changed every 48 h. The outgrowth was monitored daily and was passaged on day 4 or 5 depending on cell growth. Cultures including Tx21 were passaged into MEF-coated plates for one more passage since this facilitates attachment of the single cells. Only in passage 3 they would be converted to LT21 cultures being plated in laminin 521 pre-coated plates.

### Cell cycle analysis

After trypsinization, 10^5^ TSCs and LT21 TSCs were incubated in 0.5 ml of Tx media with 10 ug/ml Hoescht 34580 for 30 min at 37 °C. After the incubation time, tubes with cells were placed on ice and analyzed with a FACSCanto II.

### Colony formation assay

Single cells were sorted in MEF coated plates in the presence of either Tx or LT21 media. Media was changed every 48 hours and the number of wells containing colonies was assessed 7 days after sorting.

### Chimera formation

After blastocyst isolation from pregnant females, a micromanipulator was used for injecting 10 LT21 TSCs in the blastocyst cavity. Those blastocysts were then implanted in one of the uterine horns of pseudopregnant females. Each female had only one of their horns used for implantation and a maximum of 8 blastocysts were used per female. Placenta isolation was performed at E.10.5 (time of implantation was considered as 3.5). Histology was performed on the GFP imaged placentas using an antibody against GFP.

### Immunofluorescence

Samples were fixed using 4% PFA in PBS for 20 minutes at RT followed by 3 washing steps with PBS. A 0.25% triton solution in PBS was used for permeabilization during 30 minutes at RT, followed by a 1 hour blocking step with PBS+ 10% FBS and 0.1% tween20. Primary antibodies against Cdx2 (Biogenes MU392A-5UC), EOMES (Abcam ab23345), ZO-1 (Fisher scientific # 33-9100), Krt8/18 (DAKO M365201-2), GATA6 (R&D AF1700) or E-CADHERIN (life technologies # 14-3249-82) were diluted 1/200 in PBS + 0.1% Tween20, and used for staining O/N at 4C. After three washing steps, samples were incubated with the corresponding secondary antibodies for 1 hour at RT. Hoechst was used for counterstaining with or without WGA. All images were analyzed in a PerkinElmer Ultraview VoX spinning disk microscope.

### Single molecule FISH

TSCs plated in glass coverslips were allowed to grow and were then fixed using RNAse free 4%PFA in PBS + 1% Acetic Acid during 20 minutes. After fixation, all samples followed the Quantigene ViewRNA kit instructions: After three washes with RNAse free PBS, samples were incubated for 10 minutes in a detergent solution. After three washes with RNAse free PBS, samples were incubated for 5 minutes at RT with Q protease. After three washes with RNAse free PBS, samples were incubated at 40C for 3 hours (in a humidified chamber) with the probes of interest diluted in Probe set diluent. After 3 washes with wash buffer, samples were incubated at 40C for 30 minutes with preamplifier diluted in amplifier diluent. After 3 washes with wash buffer, samples were incubated at 40C for 30 minutes with amplifier diluted in amplifier diluent. After 3 washes with wash buffer, samples were incubated at 40C for 30 minutes with label diluted in label probe diluent. After 2 washes with wash buffer, they were washed once more for 10 minutes. Samples were then incubated for 15 minutes in RNAse free PBS with Hoechst and WGA as counterstains followed by 3 washes with RNAse free PBS. Blastocysts were carefully placed in mounting media in glass bottom 3.5 mm plates. All samples were imaged with a 63x oil immersion objective in a PerkinElmer Ultraview VoX spinning disk microscope.

### High content imaging

Each colony was imaged for E-CADHERIN, Cdx2, EOMES and Nuclei stainings. E-CADHERIN staining was used for manual cell segmentation in ImageJ. Cell profiler was used for analysis of cells segmentation and the other stainings. Measurements obtained in CellProfiler were used for further analysis using a Python pipeline. After discarding dividing cells based on the nuclear staining, a total of 502 control cells and 297 LT21 TSCs cells were analyzed.

### RNA sequencing

For bulk sequencing 1000 control and LT21 cultured cells were used for Trizol extraction RNA extraction from both bulk and single cells was performed following the Cel Seq 2 protocol (Hashimshony et al., 2016). Bulk samples were first normalized and then analyzed using the DESeq2 package on Rstudio. Triplicates for each group (F4 in Tx, F4 in LT21, F4 in Tx differentiated, F4 in LT21 differentiated) were analyzed. Differentially expressed were those showing an upregulation of 1.5 fold change with a p value lower than 0.05. DAVID gene ontology online tool was used for gene enrichment analysis.

### Mapping and processing of scRNA-seq data

Read 1 contains the cell or section barcode and the unique molecular identifier (UMI). Read 2 contains the biological information. Reads 2 with a valid cell barcode were selected and mapped using STAR-2.5.3a with default parameters to the mouse mm10 genome, and only reads mapping to gene bodies (exons or introns) were used for downstream analysis. Reads mapping simultaneously to an exon and to an intron were assigned to the exon. For each cell or section, the number of transcripts was obtained as previously described (Grün et al., 2014). We refer to transcripts as unique molecules based on UMI correction.

### Single molecule FISH polarity quantification

Confocal images were taken with a z-step of 0.3 μm. Given the complexity of an analysis performed in 3 dimensions that would require an algorithm capable of segmenting cells and quantifying the number of transcripts in 3D, we decided to quantify a 2 dimension projection of the slices that included the ICM and blastocoel. Those z-stack projections were oriented with polar side at the left and mural side at the right and they were then analyzed for average column pixel intensity, allowing us to plot an average pixel intensity histogram. An intensity profile was plotted for each embryo and gene. Each blastocyst is structurally different showing distinct cavity sizes, which implies that a different percentage of the TE is in contact with the ICM for each embryo. In order to compare the expression of polar and mural TE, we divided the length of the embryo in three segments of equal distance, irrespective of the total diameter. The intermediate segment was considered a transition stage between the polar and mural regions and therefore was not included in the next analyses. The polar and mural segments of the profile were analyzed by comparing the average pixel intensities of each pixel column included in the segment.

### Blastoid formation

Full protocol link: https://www.nature.com/protocolexchange-/protocols/6579). Agarose microwell arrays were casted using the PDMS stamp and incubated O/N on mES serum containing media. After washing the chips with PBS, a ESC solution of 8000 cells/150 μl is dispensed in the central chip are and allowed to settle. After 15 minutes, an additional 1ml is dispensed. 20 hours later 1 ml of media is removed and a TSC solution of 22000 cells/150 ul is dispensed. After allowing the cells to fall in the microwells, 1 ml of blastoid media is added to the wells. Blastoid media includes 20 μM Y27632 (AxonMed 1683), 3 uM CHIR99021 (AxonMed 1386), 1 mM 8Br-cAMP (Biolog Life Science Institute B007E), 25 ng/ml Fgf4 (R&D systems 5846F4), 15 ng/ml Tgfb1 (Peprotech 100-21), 30 ng/ml Il11 (Peprotech 200-11) and 1 μg/ml heparin (Sigma-Aldrich cat# H3149). An additional dose of cAMP is dispensed 24 hours after TSC seeding. All measurements are performed 65 hours after TSC seeding.

## Author Contributions

J.F.A. and N.C.R contributed to experimental designs. J.F.A. and N.C.R wrote the manuscript. J.F.A. performed all functional assays, immunostainings, smFISH, blastoid experiments, polarity experiments, and made all figures. F.D. and J.F.A performed preliminary combinatorial screens. M.K. and J.F.A. performed the factorial design-based combinatorial screen. J.V. performed CelSeq2 for all samples studied and their submission to omnibus. J.F.A and A.A.A performed the transcriptomic data analysis. L.L. performed the high-content imaging and morphometric analysis. J.K. performed blastocyst injections. A.v.O. contributed to the RNA-sequencing assays. N.G. helped supervising the project. N.C.R conceived and directed the project.

## Acknowledgements

We would like to thank Jacqueline Deschamps for providing the Cdx2-eGFP mice; Aditya Barve for preliminary data analysis. Stefan van der Elst and Reinier van der Linden for helping with FACS assays; Anka de Graaf for helping with microscopy; Harry Begthel for helping with histology; Single Cell Discoveries for the single cell transcriptomics work and all employees from the Hubrecht animal facility for their help. We sincerely thank Brian Cox for kindly discussing and correcting the manuscript.

## Accession numbers

GSE127754

---

**Figure S2.**
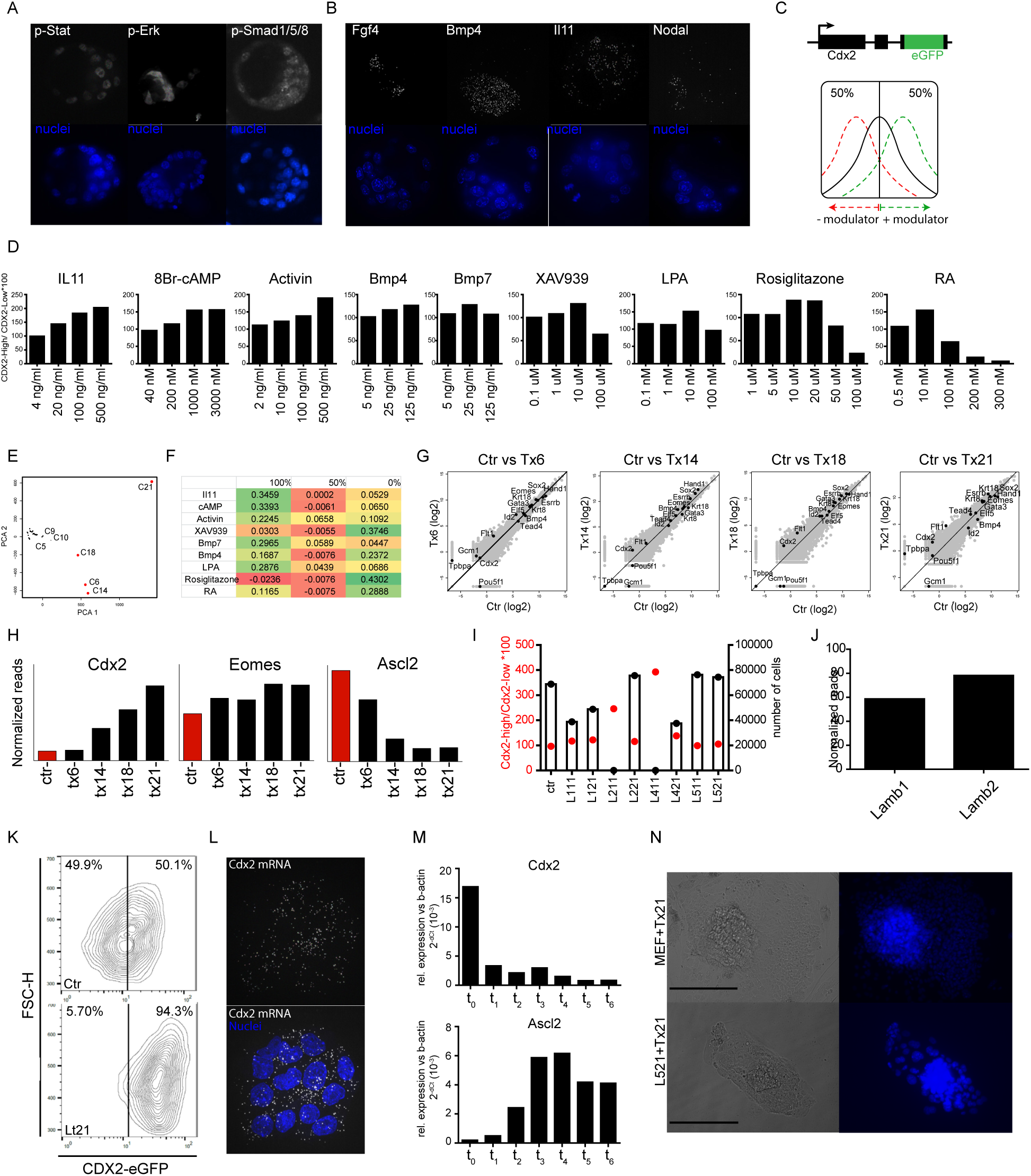
A. Pathway activation stainings performed in freshly isolated E3.5 blastocysts. B. smFISH performed on freshly isolated blastocysts allow us to detect expression of key pathway ligands. C. Cdx2 locus of CDX2-eGFP TSCs and hit detection strategy. D. Positive CDX2 expression modulators induce a higher expression of CDX2 in a dose-dependent manner. E-F. Principal component analysis suggest the effect on CDX2 expression of each compound at a given concentration. G. Bulk transcriptome analysis allows us to compare the overall expression of key trophoblast markers fr each culture condition. H. Expression of the Cdx2 and Eomes (as un-differentiated markers) and Ascl2 (as a partially differentiated marker) across the 4 best cultures. I. CDX2 expression and proliferation quantification of TSCs cultured in control conditions on plates coated with different laminins. J. Lamb1 and Lamb2 expression levels in bulk samples from blastocysts. K. Tx21 compounds combined with L521 dramatically increase the percentage of CDX2-High cells without affecting the size of the cells. L. mFISH for Cdx2 transcirpts on TSCs grown in LT21 conditions show an homogeneous expression within a colony. M. Differentiation dynamics of LT21 derived TSCs upon removal of all CDX2 regulators. A sample was taken every 24 hours after compound removal at t0. N. Proliferation comparison of trophoblasts 4 days after blastocyst plating on either mEF or L521.

**Figure S3.**
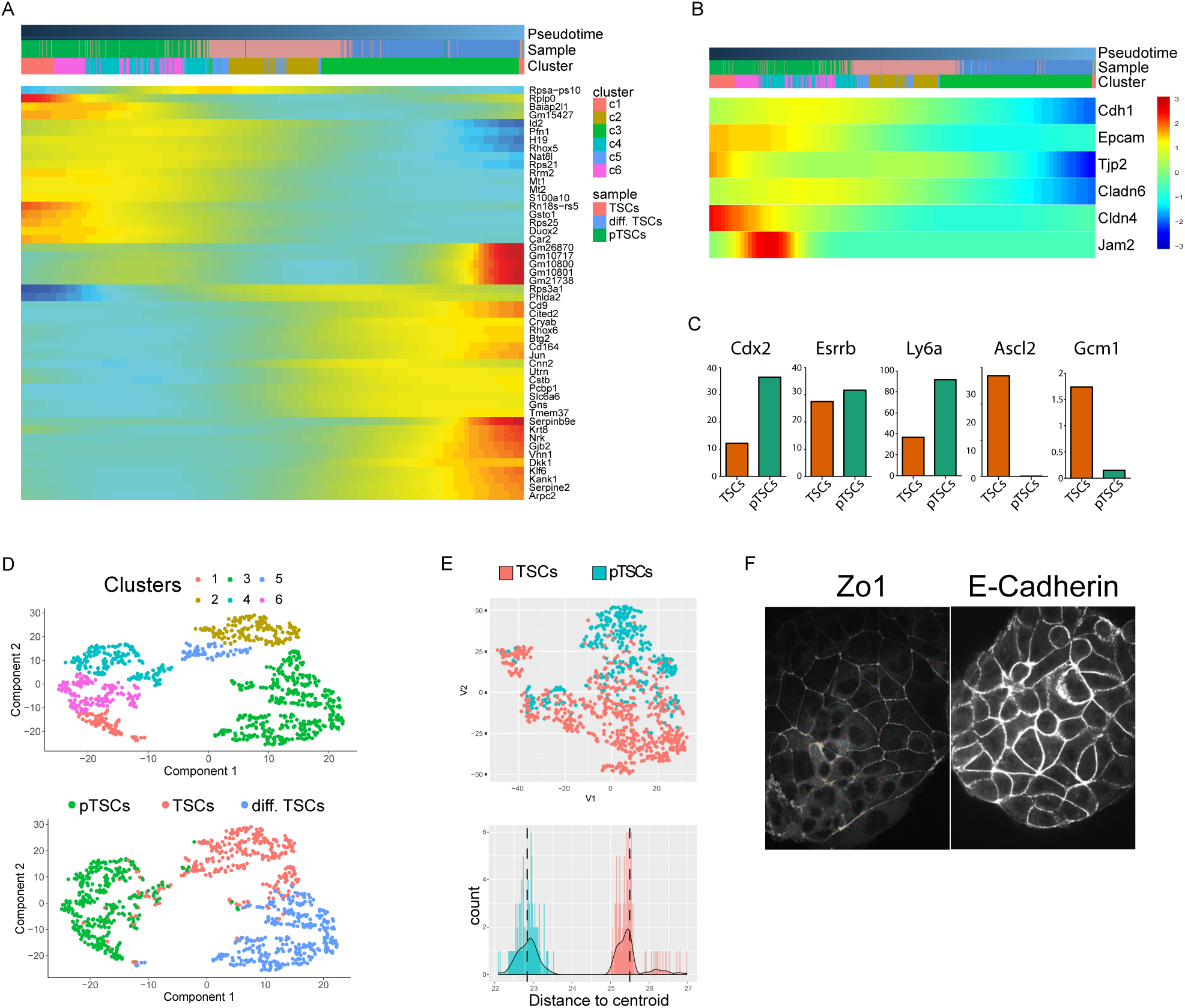
a.Top 50 genes defining the pseudotime trajectory. B. Pseudotime heatmaps for epithelial and tight junction markers. C. qPCR expression for potency and differentiation markers between control (orange) and LT21 TSCs (green). D. Monocle clustering analyisis of single cells from pTSC, TSC and differentiated TSC samples. E. Representative t-SNE map plotting each cell based on the morphological features extracted (top) and average distance to centroid for control (red) and LT21 cultured TSCs (blue). F. Expression of E-cadherin and ZO1 in LT21 cultured TSCs.

**Figure S4.**
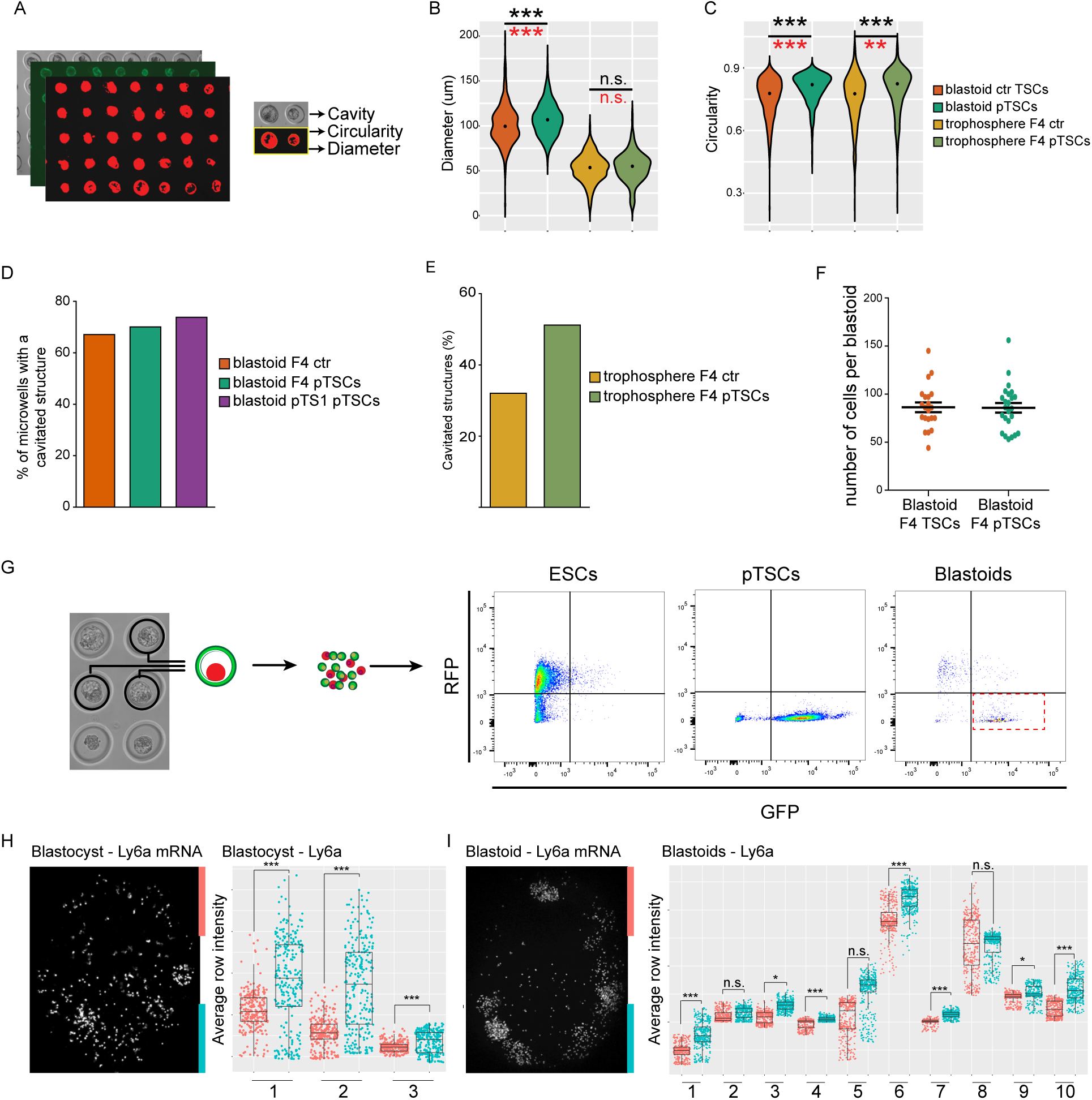
A. WGA stained structures allow us to perform a semi-automated acquisition of morphological parameters. B. Diameter of all structures for F4 TSCs and F4 pTSCs, both as blastoids and trophospheres. C. Circularity of all structures for F4 TSCs and F4 pTSCs, both as blastoids and trophospheres. D. Percentage of microwells with a cavitated structure across cultures. E. Percentage of cavitated trophospheres obtained from F4 TSCs amd and F4 pTSCs. F. Number of cells included in blastoids obtained from F4 TSCs and F4 pTSCs. G. Aproach for blastoid single trophoblast isolation. After selection of blastoids, these are digested and single GFP+ cells are sorted. H-I. smFISH staining for Ly6a on E3.5 blastocysts (H) and blastoids (I). The projection of the Z-steps including a large portion of the blastocoel was obtained and subjected to signal intensity analysis. All projections are measured in a polar-to-mural orientation. The length of the projection was divided into three segments: polar (blue), mural (red) and an intermediate segments. The latter one was not included in the analysis as it was considered a transition. The average intensity of each row of pixels is represented as a dot in the plots, which compare the intensity for rows in the polar region (blue) aginst the intensity for rows in the mural region (red) for each blastocyst (H, n=3) and blastoid measured (I, n=10). Significance assessed by two-tailed unpaired t-test. *** for pvalue < 0.001 and * for pvalue < 0.05.

**Supplementary figure 1.**
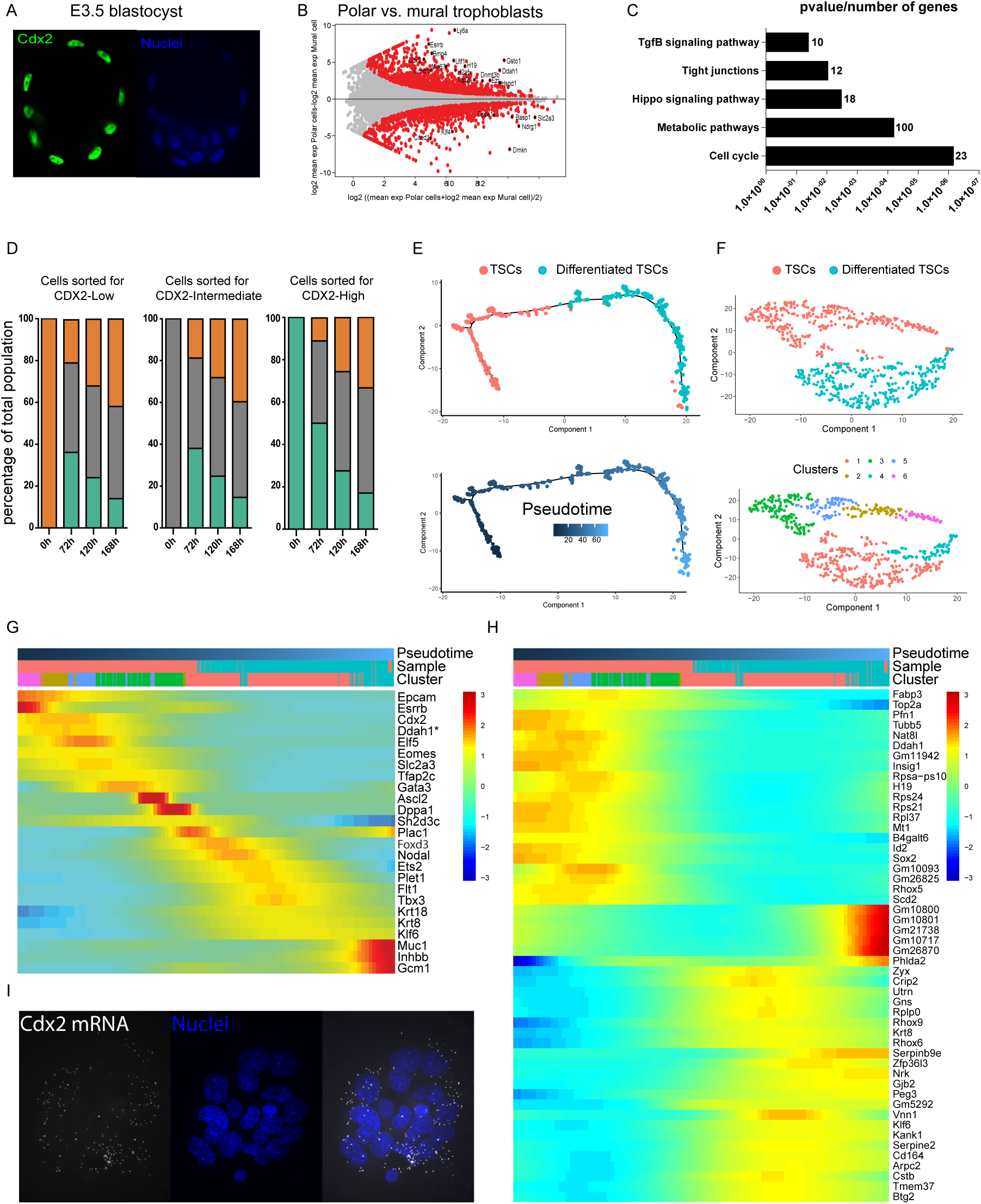
A. CDX2 staining on freshly isolated E3.5 blastocysts. B. Differential gene expression between mural and polar cells isolated from the TE of E4.5 blastocysts (Nakamura et al.). C. GO Pathway enrichement based on differential gene expression analysis of CDX2-High and CDX2-Low samples. D. Percentage of the different subpopulations upon pure subpopulation sorting and further independent culture. Green for CDX2-High, grey for CDX2-Intermediate, orange for CDX2-Low. E. Control TSCs and differentiated TSCs were located in a pseudotime trajectory by Monocle. F. Clustering analyisis of control TSCs and differentiated TSCs. G. Expression peaks for markers for varias differentiation states. Genes with * show higher expression in mural single cells as compared to polar single cells. H. Top 50 genes defining the pseudotime trajectory. I. smFISH for Cdx2 transcirpts on TSCs grown in Tx conditions show a higher expression in the outer cells of the colony. F.

## Bibliography

Auman, H.J., Nottoli, T., Lakiza, O., Winger, Q., Donaldson, S., and Williams, T. (2002). Transcription factor AP-2gamma is essential in the extra-embryonic lineages for early postimplantation development. Development 129, 2733–2747.

Baczyk, D., Satkunaratnam, A., Nait-Oumesmar, B., Huppertz, B., Cross, J.C., and Kingdom, J.C.P. (2004). Complex patterns of GCM1 mRNA and protein in villous and extravillous trophoblast cells of the human placenta. Placenta 25, 553–559.

Cao, Z., Carey, T.S., Ganguly, A., Wilson, C.A., Paul, S., and Knott, J.G. (2015). Transcription factor AP-2 induces early Cdx2 expression and represses HIPPO signaling to specify the trophectoderm lineage. Development 142, 1606–1615.

Cross, J.C., Baczyk, D., Dobric, N., Hemberger, M., Hughes, M., Simmons, D.G., Yamamoto, H., and Kingdom, J.C.P. (2003). Genes, Development and Evolution of the Placenta. Placenta 24, 123–130.

Donnison, M., Beaton, A., Davey, H.W., Broadhurst, R., L’Huillier, P., and Pfeffer, P.L. (2005). Loss of the extraembryonic ectoderm in Elf5 mutants leads to defects in embryonic patterning. Development 132, 2299–2308.

Erlebacher, A., Price, K.A., and Glimcher, L.H. (2004). Maintenance of mouse trophoblast stem cell proliferation by TGF-beta/activin. Dev. Biol. 275, 158–169.

Festuccia, N., Owens, N., and Navarro, P. (2018). Esrrb, an estrogen-related receptor involved in early development, pluripotency, and reprogramming. FEBS Lett.

Fu, Y., Wei, J., Dai, X., and Ye, Y. (2017). Increased NDRG1 expression attenuate trophoblast invasion through ERK/MMP-9 pathway in preeclampsia. Placenta 51, 76–81.

Gao, H., Gao, R., Zhang, L., Xiu, W., Zang, R., Wang, H., Zhang, Y., Gao, Y., Chen, J., and Gao, S. (2018). Esrrb plays important roles in maintaining self-renewal of trophoblast stem cells (TSCs) and reprogramming somatic cells to induced TSCs. J. Mol. Cell Biol.

Gardner, R.L. (2000). Flow of cells from polar to mural trophectoderm is polarized in the mouse blastocyst. Hum. Reprod. 15, 694–701.

Godwin, S., Ward, D., Pedone, E., Homer, M., Fletcher, A.G., and Marucci, L. (2017). An extended model for culture-dependent heterogenous gene expression and proliferation dynamics in mouse embryonic stem cells. NPJ Syst Biol Appl 3, 19.

Graham, S.J.L., and Zernicka-Goetz, M. (2016). The Acquisition of Cell Fate in Mouse Development: How Do Cells First Become Heterogeneous? Curr. Top. Dev. Biol. 117, 671–695.

Grün, D., Kester, L., and van Oudenaarden, A. (2014). Validation of noise models for single-cell transcriptomics. Nat. Methods 11, 637–640.

Guillemot, F., Nagy, A., Auerbach, A., Rossant, J., and Joyner, A.L. (1994). Essential role of Mash-2 in extraembryonic development. Nature 371, 333–336.

Guzman-Ayala, M., Ben-Haim, N., Beck, S., and Constam, D.B. (2004). Nodal protein processing and fibroblast growth factor 4 synergize to maintain a trophoblast stem cell microenvironment. Proc. Natl. Acad. Sci. U. S. A. 101, 15656–15660.

Hanna, L.A., Foreman, R.K., Tarasenko, I.A., Kessler, D.S., and Labosky, P.A. (2002). Requirement for Foxd3 in maintaining pluripotent cells of the early mouse embryo. Genes Dev. 16, 2650–2661.

Hastreiter, S., and Schroeder, T. (2016). Nanog dynamics in single embryonic stem cells. Cell Cycle 15, 770–771.

Hemberger, M., Udayashankar, R., Tesar, P., Moore, H., and Burton, G.J. (2010). ELF5-enforced transcriptional networks define an epigenetically regulated trophoblast stem cell compartment in the human placenta. Hum. Mol. Genet. 19, 2456–2467.

Hughes, M., Dobric, N., Scott, I.C., Su, L., Starovic, M., St-Pierre, B., Egan, S.E., Kingdom, J.C.P., and Cross, J.C. (2004). The Hand1, Stra13 and Gcm1 transcription factors override FGF signaling to promote terminal differentiation of trophoblast stem cells. Dev. Biol. 271, 26–37.

Hutchens, S.A., León, R.V., O’neill, H.M., and Evans, B.R. (2007). Statistical analysis of optimal culture conditions for Gluconacetobacter hansenii cellulose production. Lett. Appl. Microbiol. 44, 175–180.

Imakawa, K., Dhakal, P., Kubota, K., Kusama, K., Chakraborty, D., Karim Rumi, M.A., and Soares, M.J. (2016). CITED2 modulation of trophoblast cell differentiation: insights from global transcriptome analysis. Reproduction 151, 509–516.

Kidder, B.L., and Palmer, S. (2010). Examination of transcriptional networks reveals an important role for TCFAP2C, SMARCA4, and EOMES in trophoblast stem cell maintenance. Genome Res. 20, 458–472.

Kuales, G., Weiss, M., Sedelmeier, O., Pfeifer, D., and Arnold, S.J. (2015). A Resource for the Transcriptional Signature of Bona Fide Trophoblast Stem Cells and Analysis of Their Embryonic Persistence. Stem Cells Int. 2015, 218518.

Kubaczka, C., Senner, C., Araúzo-Bravo, M.J., Sharma, N., Kuckenberg, P., Becker, A., Zimmer, A., Brüstle, O., Peitz, M., Hemberger, M., et al. (2014). Derivation and maintenance of murine trophoblast stem cells under defined conditions. Stem Cell Reports 2, 232–242.

Kuckenberg, P., Buhl, S., Woynecki, T., van Fürden, B., Tolkunova, E., Seiffe, F., Moser, M., Tomilin, A., Winterhager, E., and Schorle, H. (2010). The transcription factor TCFAP2C/AP-2gamma cooperates with CDX2 to maintain trophectoderm formation. Mol. Cell. Biol. 30, 3310–3320.

Larue, L., Ohsugi, M., Hirchenhain, J., and Kemler, R. (1994). E-cadherin null mutant embryos fail to form a trophectoderm epithelium. Proceedings of the National Academy of Sciences 91, 8263–8267.

Latos, P.A., Goncalves, A., Oxley, D., Mohammed, H., Turro, E., and Hemberger, M. (2015a). Fgf and Esrrb integrate epigenetic and transcriptional networks that regulate self-renewal of trophoblast stem cells. Nat. Commun. 6, 7776.

Latos, P.A., Sienerth, A.R., Murray, A., Senner, C.E., Muto, M., Ikawa, M., Oxley, D., Burge, S., Cox, B.J., and Hemberger, M. (2015b). Elf5-centered transcription factor hub controls trophoblast stem cell self-renewal and differentiation through stoichiometry-sensitive shifts in target gene networks. Genes Dev. 29, 2435–2448.

Lin, S.-C.J., Wani, M.A., Whitsett, J.A., and Wells, J.M. (2010). Klf5 regulates lineage formation in the pre-implantation mouse embryo. Development 137, 3953–3963.

Luo, J., Sladek, R., Bader, J.-A., Matthyssen, A., Rossant, J., and Giguère, V. (1997). Placental abnormalities in mouse embryos lacking the orphan nuclear receptor ERR-β. Nature 388, 778–782.

Luo, Y., Lim, C.L., Nichols, J., Martinez-Arias, A., and Wernisch, L. (2013). Cell signalling regulates dynamics of Nanog distribution in embryonic stem cell populations. J. R. Soc. Interface 10, 20120525.

Ma, G.T., Roth, M.E., Groskopf, J.C., Tsai, F.Y., Orkin, S.H., Grosveld, F., Engel, J.D., and Linzer, D.I. (1997). GATA-2 and GATA-3 regulate trophoblast-specific gene expression in vivo. Development 124, 907–914.

McDole, K., and Zheng, Y. (2012). Generation and live imaging of an endogenous Cdx2 reporter mouse line. Genesis 50, 775–782.

Menchero, S., Rayon, T., Andreu, M.J., and Manzanares, M. (2017). Signaling pathways in mammalian preimplantation development: Linking cellular phenotypes to lineage decisions. Dev. Dyn. 246, 245–261.

Morioka, Y., Nam, J.-M., and Ohashi, T. (2017). Nik-related kinase regulates trophoblast proliferation and placental development by modulating AKT phosphorylation. PLoS One 12, e0171503.

Moriwaki, K., Tsukita, S., and Furuse, M. (2007). Tight junctions containing claudin 4 and 6 are essential for blastocyst formation in preimplantation mouse embryos. Dev. Biol. 312, 509–522.

Motomura, K., Oikawa, M., Hirose, M., Honda, A., Togayachi, S., Miyoshi, H., Ohinata, Y., Sugimoto, M., Abe, K., Inoue, K., et al. (2016). Cellular Dynamics of Mouse Trophoblast Stem Cells: Identification of a Persistent Stem Cell Type. Biol. Reprod. 94, 122.

Nakamura, T., Yabuta, Y., Okamoto, I., Aramaki, S., Yokobayashi, S., Kurimoto, K., Sekiguchi, K., Nakagawa, M., Yamamoto, T., and Saitou, M. (2015). SC3-seq: a method for highly parallel and quantitative measurement of single-cell gene expression. Nucleic Acids Res. 43, e60.

Natale, B.V., Schweitzer, C., Hughes, M., Globisch, M.A., Kotadia, R., Tremblay, E., Vu, P., Cross, J.C., and Natale, D.R.C. (2017). Sca-1 identifies a trophoblast population with multipotent potential in the mid-gestation mouse placenta. Sci. Rep. 7, 5575.

Natale, D.R.C., Hemberger, M., Hughes, M., and Cross, J.C. (2009). Activin promotes differentiation of cultured mouse trophoblast stem cells towards a labyrinth cell fate. Dev. Biol. 335, 120–131.

Ng, R.K., Dean, W., Dawson, C., Lucifero, D., Madeja, Z., Reik, W., and Hemberger, M. (2008). Epigenetic restriction of embryonic cell lineage fate by methylation of Elf5. Nat. Cell Biol. 10, 1280–1290.

Niakan, K.K., Schrode, N., Cho, L.T.Y., and Hadjantonakis, A.-K. (2013). Derivation of extraembryonic endoderm stem (XEN) cells from mouse embryos and embryonic stem cells. Nat. Protoc. 8, 1028–1041.

Nishioka, N., Yamamoto, S., Kiyonari, H., Sato, H., Sawada, A., Ota, M., Nakao, K., and Sasaki, H. (2008). Tead4 is required for specification of trophectoderm in pre-implantation mouse embryos. Mech. Dev. 125, 270–283.

Nishioka, N., Inoue, K.-I., Adachi, K., Kiyonari, H., Ota, M., Ralston, A., Yabuta, N., Hirahara, S., Stephenson, R.O., Ogonuki, N., et al. (2009). The Hippo signaling pathway components Lats and Yap pattern Tead4 activity to distinguish mouse trophectoderm from inner cell mass. Dev. Cell 16, 398–410.

Niwa, H., Toyooka, Y., Shimosato, D., Strumpf, D., Takahashi, K., Yagi, R., and Rossant, J. (2005). Interaction between Oct3/4 and Cdx2 determines trophectoderm differentiation. Cell 123, 917–929.

Ochiai, H., Sugawara, T., Sakuma, T., and Yamamoto, T. (2014). Stochastic promoter activation affects Nanog expression variability in mouse embryonic stem cells. Sci. Rep. 4, 7125.

Ohinata, Y., and Tsukiyama, T. (2014). Establishment of trophoblast stem cells under defined culture conditions in mice. PLoS One 9, e107308.

Paul, S., and Home, P. (2015). Combinatorial functions of GATA2 and GATA3 are essential for early trophoblast development and to balance the stem vs. differentiation and angiogenic equilibrium in trophoblast lineage. Placenta 36, A34.

Posfai, E., Petropoulos, S., de Barros, F.R.O., Schell, J.P., Jurisica, I., Sandberg, R., Lanner, F., and Rossant, J. (2017). Position- and Hippo signaling-dependent plasticity during lineage segregation in the early mouse embryo. Elife 6.

Ralston, A., Cox, B.J., Nishioka, N., Sasaki, H., Chea, E., Rugg-Gunn, P., Guo, G., Robson, P., Draper, J.S., and Rossant, J. (2010). Gata3 regulates trophoblast development downstream of Tead4 and in parallel to Cdx2. Development 137, 395–403.

Ray, S., Dutta, D., Karim Rumi, M.A., Kent, L.N., Soares, M.J., and Paul, S. (2008). Context-dependent Function of Regulatory Elements and a Switch in Chromatin Occupancy between GATA3 and GATA2 RegulateGata2Transcription during Trophoblast Differentiation. J. Biol. Chem. 284, 4978–4988.

Rayon, T., Menchero, S., Nieto, A., Xenopoulos, P., Crespo, M., Cockburn, K., Cañon, S., Sasaki, H., Hadjantonakis, A.-K., de la Pompa, J.L., et al. (2014). Notch and hippo converge on Cdx2 to specify the trophectoderm lineage in the mouse blastocyst. Dev. Cell 30, 410–422.

Rivron, N. (2018). Formation of blastoids from mouse embryonic and trophoblast stem cells.

Rivron, N., Pera, M., Rossant, J., Martinez Arias, A., Zernicka-Goetz, M., Fu, J., van den Brink, S., Bredenoord, A., Dondorp, W., de Wert, G., et al. (2018a). Debate ethics of embryo models from stem cells. Nature 564, 183–185.

Rivron, N.C., Frias-Aldeguer, J., Vrij, E.J., Boisset, J.-C., Korving, J., Vivié, J., Truckenmüller, R.K., van Oudenaarden, A., van Blitterswijk, C.A., and Geijsen, N. (2018b). Blastocyst-like structures generated solely from stem cells. Nature 557, 106–111.

Roberts, R.M., Michael Roberts, R., and Fisher, S.J. (2011). Trophoblast Stem Cells1. Biol. Reprod. 84, 412–421.

Rossant, J. (2001). Stem Cells from the Mammalian Blastocyst. Stem Cells 19, 477–482.

Rossant, J. (2004). Lineage development and polar asymmetries in the peri-implantation mouse blastocyst. Semin. Cell Dev. Biol. 15, 573–581.

Rossant, J., and Tam, P.P.L. (2009). Blastocyst lineage formation, early embryonic asymmetries and axis patterning in the mouse. Development 136, 701–713.

Russ, A.P., Wattler, S., Colledge, W.H., Aparicio, S.A., Carlton, M.B., Pearce, J.J., Barton, S.C., Surani, M.A., Ryan, K., Nehls, M.C., et al. (2000). Eomesodermin is required for mouse trophoblast development and mesoderm formation. Nature 404, 95–99.

Schmidt, S., Hommel, A., Gawlik, V., Augustin, R., Junicke, N., Florian, S., Richter, M., Walther, D.J., Montag, D., Joost, H.-G., et al. (2009). Essential role of glucose transporter GLUT3 for post-implantation embryonic development. J. Endocrinol. 200, 23–33.

Shahbazi, M.N., and Zernicka-Goetz, M. (2018). Deconstructing and reconstructing the mouse and human early embryo. Nat. Cell Biol. 20, 878–887.

Shi, X.-H., Larkin, J.C., Chen, B., and Sadovsky, Y. (2013). The expression and localization of N-myc downstream-regulated gene 1 in human trophoblasts. PLoS One 8, e75473.

Simmons, D.G., and Cross, J.C. (2005). Determinants of trophoblast lineage and cell subtype specification in the mouse placenta. Dev. Biol. 284, 12–24.

Sood, R., Kalloway, S., Mast, A.E., Hillard, C.J., and Weiler, H. (2006a). Fetomaternal cross talk in the placental vascular bed: control of coagulation by trophoblast cells. Blood 107, 3173–3180.

Sood, R., Zehnder, J.L., Druzin, M.L., and Brown, P.O. (2006b). Gene expression patterns in human placenta. Proc. Natl. Acad. Sci. U. S. A. 103, 5478–5483.

Strumpf, D., Mao, C.-A., Yamanaka, Y., Ralston, A., Chawengsaksophak, K., Beck, F., and Rossant, J. (2005). Cdx2 is required for correct cell fate specification and differentiation of trophectoderm in the mouse blastocyst. Development 132, 2093–2102.

Tanaka, S., Kunath, T., Hadjantonakis, A.K., Nagy, A., and Rossant, J. (1998). Promotion of trophoblast stem cell proliferation by FGF4. Science 282, 2072–2075.

Tompers, D.M., Foreman, R.K., Wang, Q., Kumanova, M., and Labosky, P.A. (2005). Foxd3 is required in the trophoblast progenitor cell lineage of the mouse embryo. Dev. Biol. 285, 126–137.

Trapnell, C., Cacchiarelli, D., Grimsby, J., Pokharel, P., Li, S., Morse, M., Lennon, N.J., Livak, K.J., Mikkelsen, T.S., and Rinn, J.L. (2014). The dynamics and regulators of cell fate decisions are revealed by pseudotemporal ordering of single cells. Nat. Biotechnol. 32, 381–386.

Vrij, E.J., Reimer, Y.S.O., Aldeguer, J.F., Guerreiro, I.M., Kind, J., Koo, B.-K., van Blitterswijk, C., and Rivron, N. (2019). Self-organization of post-implantation-like embryonic tissues from blastoids.

Wen, F., Tynan, J.A., Cecena, G., Williams, R., Múnera, J., Mavrothalassitis, G., and Oshima, R.G. (2007). Ets2 is required for trophoblast stem cell self-renewal. Dev. Biol. 312, 284–299.

Yagi, R., Kohn, M.J., Karavanova, I., Kaneko, K.J., Vullhorst, D., DePamphilis, M.L., and Buonanno, A. (2007). Transcription factor TEAD4 specifies the trophectoderm lineage at the beginning of mammalian development. Development 134, 3827–3836.

Ying, Q.-L., Wray, J., Nichols, J., Batlle-Morera, L., Doble, B., Woodgett, J., Cohen, P., and Smith, A. (2008). The ground state of embryonic stem cell self-renewal. Nature 453, 519–523.

Zhang, H.T., and Hiiragi, T. (2018). Symmetry Breaking in the Mammalian Embryo. Annu. Rev. Cell Dev. Biol. 34, 405–426.

